# Comprehensive Analysis of Co-Mutations Identifies Cooperating Mechanisms of Tumorigenesis

**DOI:** 10.1101/2021.08.23.457315

**Authors:** Limin Jiang, Hui Yu, Scott Ness, Peng Mao, Fei Guo, Jijun Tang, Yan Guo

## Abstract

Somatic mutations are one of the most important factors in tumorigenesis and are the focus of most cancer sequencing efforts. The co-occurrence of multiple mutations in one tumor has gained increasing attention as a means of identifying cooperating mutations or pathways that contribute to cancer.Using multi-omics, phenotypical, and clinical data from 29,559 cancer subjects and 1,747 cancer cell lines covering 78 distinct cancer types, we show that co-mutations are associated with prognosis, drug sensitivity, and disparities in sex, age, and race. Some co-mutation combinations displayed stronger effects than their corresponding single mutations. For example, co-mutation *TP53*:*KRAS* in pancreatic adenocarcinoma is significantly associated with disease specific survival (hazard ratio = 2.87, adjusted p-value = 0.0003) and its prognostic predictive power is greater than either *TP53* or *KRAS* as individually mutated genes. Functional analyses revealed that co-mutations with higher prognostic values have higher potential impact and cause greater dysregulation of gene expression. Furthermore, many of the prognostically significant co-mutations caused gains or losses of binding sequences of RNA binding proteins or micro RNAs with known cancer associations. Thus, detailed analyses of co-mutations can identify mechanisms that cooperate in tumorigenesis.

## Introduction

Tumors acquire somatic mutations in oncogenes and tumor suppressors that lead to tumorigenesis [1]. While most studies of somatic mutations focus on the impact of single mutations, researchers have started to appreciate the cooperative effects induced by multiple mutations arising simultaneously in one tumor. The event of multiple mutations emerging concurrently is referred to as co-mutation or mutation co-occurrence. Because genes are the basic genomic unit that bears a more-or-less self-contained function, researchers usually identify mutated genes and study the co-mutations between two (or multiple) distinct genes. Many studies have suggested that co-mutation is a core determinant of oncogene-driven cancers. For example, co-mutations have been shown to be associated with pathogenesis, immune microenvironment, therapeutic vulnerabilities of cancer, and drug sensitivity in non-small-cell lung cancer (NSCLC) [2]. Lung cancer patients with co-mutation of *EGFR, TP53*, and *RB1* have a higher risk of histologic transformation [3]. Co-mutation is also a major determinant of the molecular diversity of *KRAS*-mutant lung adenocarcinomas [4]. *TET2*–*SRSF2* co-mutation has a strong association with the chronic myelomonocytic leukemia phenotype - the larger the *TET2–SRSF2* co-mutated clone, the more prominent the monocytosis [5]. *ARID1A*:*PIK3CA* co-mutation in the endometrial epithelium promotes an invasive phenotype [6].

A number of studies have revealed associations between co-mutations and clinical outcomes. For example, *TP53*:*KRAS* co-mutation in NSCLC was found to confer clinical benefit to PD-1 inhibitors [7]. *CREBBP*:*STAT6* co-mutation supports the diagnosis of the diffuse variant of follicular lymphoma [8]. NSCLC patients with *EGFR:TP53 or EGFR:PIK3CA* co-mutation are more likely to be resistant to the first-generation *EGFR* tyrosine kinase inhibitors [9]. In general, co-mutation demonstrates a prognostic value in vulvar squamous cell carcinoma (VSCC) [10], NSCLC [11], acute myeloid leukemia (AML) [12], and lung adenocarcinoma (LUAD) [13].

Previous co-mutation studies were generally conducted focusing on individual cancer types and have not systematically interrogated all combinations of protein-coding genes and non-coding genes. In the present work, we performed a comprehensive pan-cancer co-mutation study that integrated multi-omics data from ∼30,000 subjects of over 50 cancer types from diverse cancer consortiums. We set our analysis perspective both at nucleotide base level and gene level, and extended the co-mutation search scope to the full domain of protein-coding genes and non-coding genes. Functional associations of co-mutation instances with cancer prognosis, cis-regulatory elements, and transcription dysregulations were also thoroughly examined. The results support previous models of oncogene cooperativity and the multi-hit hypothesis, but also identify new types of cooperation between important genes involved in tumorigenesis.

## Methods

### Data acquisition

Somatic mutation data and gene expression data (RNA-Seq FPKM) of 10,147 TCGA subjects were downloaded from the Genomic Data Commons. We used TCGA Pan-Cancer Clinical Data Resource [14] to acquire disease specific survival information. ICGC mutation and clinical data of 19,412 subjects were downloaded from ICGC dataportal. Mutation and gene expression of 1,747 cancer cell lines were download from DepMap, previously known as the Cancer Cell Line Encyclopedia (CCLE). The drug sensitivity data of 4,686 drugs were also downloaded from DepMap. Some phenotypical variables were available and downloaded (TCGA: age, sex, and race; ICGC: age and sex). TCGA is a consortium that originated in the US, all subjects were recruited in US. ICGC in an international consortium, it contains 57 cancer types from 81 cohorts. Some cohorts share the same cancer type. There may be a small portion of overlapping data between the ICGC and TCGA. Because we performed separated analyses of ICGC and TCGA, we did not attempt to identify the overlapping subjects. The numerous subjects or cell lines were grouped by cancer type (TCGA), cohort (ICGC), or tissue site (DepMap), and we excluded any dataset with sample size ≤ 50.

### Mutation annotation

All types of mutations, including single-nucleotide substitutions, insertions, and deletions, were covered in our analysis. We used ANNOVAR [15] to characterize regional and functional categories for each genomic mutation that was located to an accurate chromosome coordinate position. Gene types (protein-coding and non-coding) were derived from the latest GENCODE gene transfer format (GTF) file v34. As a common practice, we dropped the synonymous mutations from the protein-coding mutation set because of their negligible influence on protein sequences. When a quantity of gene length was necessary for an analysis, we calculated the distance between the transcription start site and the transcription end site. In describing the circumstances of single mutations (as opposed to co-mutations), we defined a mutation frequency with respect to a cohort as the fraction of subjects carrying the mutation in question. At times, we may talk about mutation frequency at the gene level, in which context we referred to the fraction of subjects having at least one mutation in the concerned gene.

### Co-mutation definition

Co-mutation was classified at two different levels: gene level and position level. At position level, the exact genomic position displaying a mutation was considered a unique entity and two positions bearing mutations in a same genome (same subject) formed a co-mutation pair. At gene level, two genes were deemed as a co-mutation pair as long as any cross-gene concurrent mutations appeared; the actual number of cross-gene co-mutation instances were not taken into account. For example, if one sample harbors two mutations in gene A and three mutations in gene B, we consider only one co-mutation pair (Gene A:Gene B) at the gene level, but six (i.e., 2×3) co-mutation pairs at position level. A co-mutation pair was supported by a quantitative metric of frequency, defined as the fraction of subjects harboring concurrent mutations in the concerned entity pair. Throughout this work, we only analyzed co-mutation pairs of frequencies ⩾ 10%. Because genes can be divided into a protein-coding set and a non-coding set, we studied three types of co-mutation gene pairs: coding:coding, coding:non-coding, and non-coding:non-coding. Finally, based on the discrete chromosomes, we differentiated co-mutation pairs into inter-chromosome ones and intra-chromosome ones. Co-mutated gene pairs that were located on one same chromosome were designated as intra-chromosome pairs, and the co-mutated gene pairs that each involved two distinct chromosomes were designated as inter-chromosome pairs.

### Phenotypic variable association Analysis

We conducted association analysis between each co-mutation gene pair and each phenotypic variable. Each subject was asigned a binary value (0 or 1) for the co-mutation variable, which designated whether or not the two genes were both mutated in the subject. Additionally, each subject was asigned a binary, multi-nomial, or continuous value for the phenotypic variable, depending on its nature. Within the scope of a subject group (cohort, cancer type, or tissue site), multiple subjects contributed values for the dependent variable (co-mutation) and the response variable (phenotype), and thereby allowed us to screen for co-mutation gene pairs that were significantly associated with a phenotypic variable. Because of the varied natures, the age variable used linear regression, the sex variable used logistic regression, and the race variable used multi-nominal regression. In the analysis for the sex variable, we coded 1 for male and 0 for female, and did not analyze gender-specific cancers such as breast cancer and prostate cancer.

### Survival Analysis

We conducted survival analysis for each co-mutation gene pair within each cancer cohort, in largely the same way as we did in the phenotype association analyses. The binary co-mutation variable denoted if a subject harbored the concurrent mutations or not, and the prognosis prediction ability of the co-mutation was assessed with Cox proportional hazard regression model. Prognosis information came in the form of disease speficic survival for TCGA and overall survival for ICGC. Multiple test correction was peformed with the Benjamini-Hochberg method. An adjusted p-value less than 0.05 was considered statistically significant, and an adjusted p-value in the interval of [0.05, 0.1] was considered marginally significant. During survival analysis, there is a chance that all events were allocated to either the mutant or the wildtype group. In such a scenario, the Cox proportional hazard model will not converge, the hazard ratio (HR) reported would be infinity. Thus, in the scenario where one of the groups (mutant and wildtype) did not receive any events, we simply asserted the co-mutation as significantly associated with survival due to imblalanced events. As a result, the returned prognostic co-mutations were ascertained with three different levels of significance: 1) empirical significance due to imbalanced events; 2) significance with adjusted p-value < 0.05; 3) marginal significance with adjusted p-value falling in [0.05, 0.1].

Mutational burden generally refers to the total amount of mutations across a single human genome, which is found an informative aggregate index in cancer biology. Henceforth, we also conducted survival analyses with mutational burden of a single mutation or a co-mutation as the dependent variable. Adjusted p-value < 0.05 was used as the significant threshold. To demonstrate that co-mutation’s prognostic value is not a byproduct of single mutations, we performed survival between mutant and wildtype groups based on single mutations and compared the results between single mutation and co-mutation.

### Regulatory element analysis

When mutation takes place in cis-regulatory elements, regulation of gene expression may be affected and the impact of a mutation may be propagated to a large number of regulatory targets [16]. We leveraged Somatic Binding Sequence Analyzer [17] to identify cis-regulatory elements affected by each mutation of a co-mutation pair. Technically, we screened three classes of cis-regulatory elements, namely RNA-binding protein (RBP) binding sequences, miRNA seed sequences, and miRNA-matching 3’-UTR sequences. RBP binding sequences numbered 3,524 and were downloaded from four databases: ATtRACT [18], ORNAment [19], RBPDB [20], and RBPmap [21]. MiRNA seed sequences numbered 2,879 and were downloaded from mirBase [22]. MiRNA-matching 3’-UTR sequences numbered 2,055,403 and were downloaded from starBase 2.0 [23]. Circos plot [24] was used to manifest a genome-wide view of affected cis-regulatory elements.

### Mutation impact analysis

A series of methods are available to assess the functional impact resulting from a mutation at a particular genomic position. These methods are generally based on multiple sequence alignment within a protein family, presuming that positions with a low conservation rate are likely to tolerate a mutation while positions with a high conversion rate are likely to be intolerant to a mutation. In light of such a conversational perspective, mutation impact was predicted for each genomic position of each co-mutation gene pair, using eight algorithms: SIFT[25], Polyphen2 (including both HDIV and HVAR) [26], LRT [27], FATHMM [28], CADD [29], VEST3 [30], and MetaSVM [31]. The scores out of distinct algorithms were normalized to a common scale between 0 to 1, where a higher value signified a stronger impact. To summarize the postion-level impact scores to the gene level, an average impact score was obtained across all mutated positions for the co-mutation gene pair in question. For each gene level co-mutation, the mutation impact is the average prediction algorithm score of all point mutations within the two genes from this co-mutation.

In addition to these theory based methods, we also utilized several empirical data based methods. Drug sensitivity difference between co-mutation mutant and wildtype groups were conducted using t-test, where an adjust p-value < 0.05 was considered statistical significant. Furthermore, a previous study [32] showed that mutations with high impact tend to cause more gene expression dysregulation. Based on this concept, we examined the differential gene expression between co-mutation mutatant and wildtype groups.

### Clinical cancer gene panels

Four panels of clinically relevant cancer genes were commonly leveraged in cancer researches, namely Agilent SureSelect (98 genes), University of California San Francisco UCSF500 (529 genes), FoundationOne CDx (309 genes), and Ashion Genomic Enabled Medicine (540 genes). Genes harboring prognostic co-mutations were compared against these four clinical cancer gene panels using R package UpSetR [33].

## Results

### Overall single mutation description

The three major data sources underlying our study, TCGA, ICGC, and DepMap, were organized in terms of cancer types, cohorts, and tissue sites. Before we moved on to the central topic of co-mutation, we first gave the data a comprehensive description from the perspective of single mutations. For each subject, we counted the number of genes bearing at least one mutation; the numbers of mutated genes per subject were displayed to reveal disparity across cancers (Figure 1). Because DNA mismatch repair genes (*MLH1, MLH3, MSH3, MSH6, PMS1, PMS2*, and *PMS2L3*) and *POLE* are frequently associated with hypermutation [34], we distinguished subjects bearing mutations in these genes. Several interesting phenomena came to our attention. First, as expected, cancer types with higher mutational loads also included more subjects having mutations in DNA mismatch repair genes or *POLE*. This observation reiterates the effect of mutations of DNA mismatch repair genes or *POLE* on the overall mutational burden. Additionally, evident hypermutation groups were observed in several cancer types. The most conspicuous hypermutation group existed in TCGA’s uterine corpus endometrial carcinoma (UCEC) cohort (Figure 1A), which seemed to be predominated by subjects having both mutated DNA mismatch repair genes and mutated *POLE*. A similar hypermutation group can be seen in the counterpart cohort in ICGC, UCEC-US (Figure 1B). The hypermutation phenomenon is closely related to several characteristics we observed for co-mutations in the UCEC cohort.

**Figure 1.**
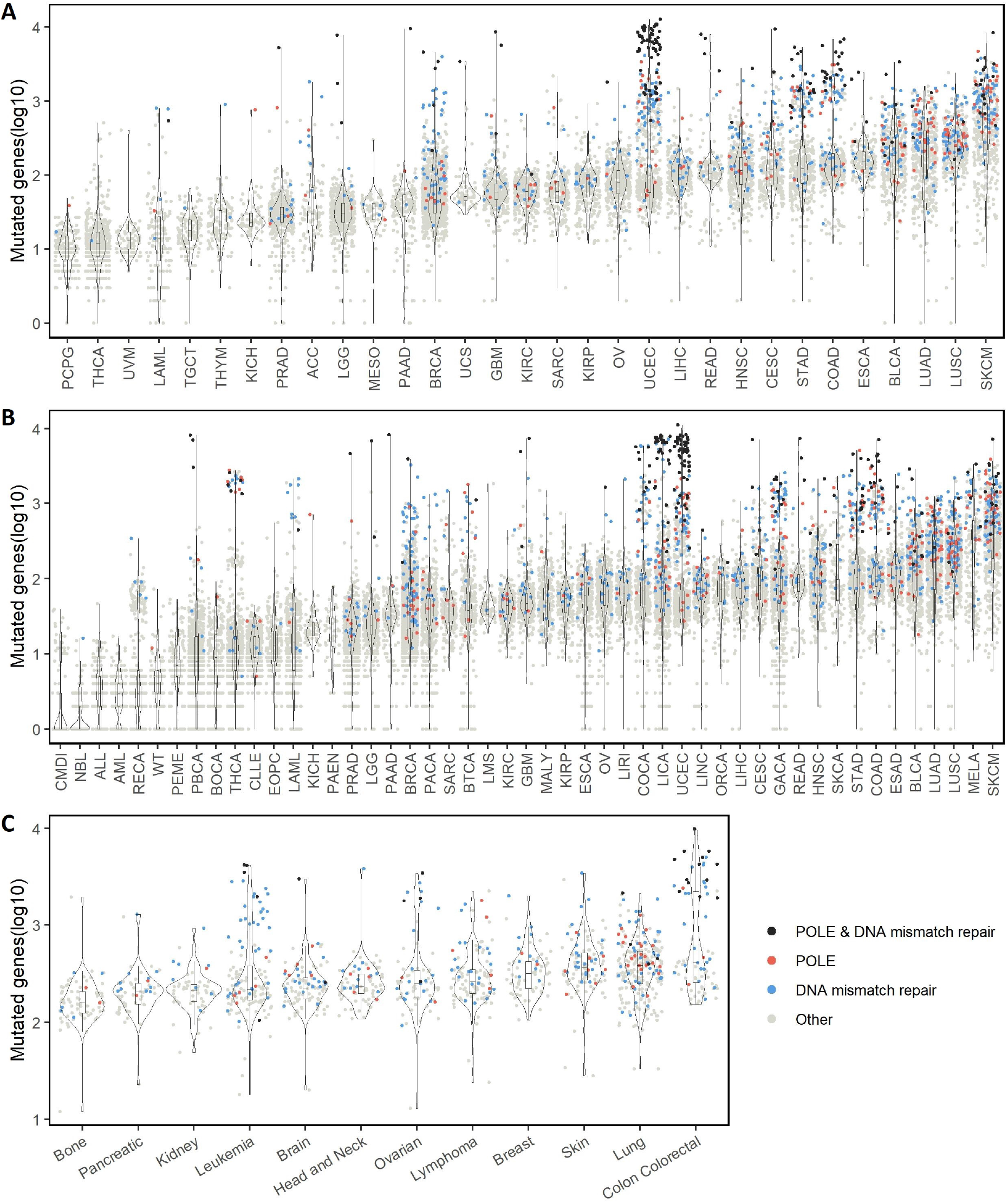
Number of mutated genes per subject of each cancer cohort. Results are reported for three cancer data consortiums separately: A. The Cancer Genome Atlas (TCGA). B. International Cancer Genome Consortium (ICGC). C. Cancer Dependency Map (DepMap). Only cohorts with sample size ⩾ 50 were drawn. Each data-point represents one genome sample (subject or cell line). Dot color signifies mutation in a specific category of genes: blue, DNA mismatch repair genes; red, gene *POLE*; black, DNA mismatch repair genes as well as gene *POLE*; gray, all other genes.

Certain cancer types showed a distinctive bimodality in the distribution of per-subject mutated genes. Using Hartigan’s Dip Test of Unimodality, five TCGA cohorts, colon adenocarcinoma (COAD), acute myeloid leukemia (LAML), pheochromocytoma and paraganglioma (PCPG), thyroid carcinoma (THCA), and UCEC were found with significant multimodality (FDR-adjusted p-value < 0.05). For example, among the 404 patients of TCGA’s COAD cohort, where 80% of the patients had less than 300 mutated genes, 18% of patients had more than 700 mutated genes, leaving a visible gap between the groups. The bimodality in the mutated gene quantity distribution suggests multiple different mechanisms are likely to be responsible for cancer formation and development. Somatic mutations may be the primary tumorigenesis cause amongst patients with a large number of mutated genes; for patients with a low number of mutated genes other genomic aberrations such as copy number variation or post transcriptional modification may have a major impact [35].

The three sources of data did not use a same technology to capture mutations. TCGA used exome sequencing, ICGC used whole genome sequencing, DepMap used whole genome sequencing but released only exonic mutations as of this writing. The numbers of mutations generated from each consortium were rather different. The average numbers of mutations per subject/cell line were 276, 170, and 507 for TCGA, ICGC and DepMap, respectively. Noticeably, DepMap had a much greater number of mutations per sample than either TCGA or ICGC, which may be a reflection of the distinct nature of cell lines. Tumor samples are usually the combination of tumor and normal cells, while the tumor cell lines are a pure clone originating from a single origin tumor cell. Thus, mutations are easier to detect in cell lines than in tumors. In addition, cell lines have been selected for growth in culture, which could select for acdditional mutations. In DepMap the tissue site with the greatest number of mutated genes is colon, where a bimodality distribution was noticeable as in the colon cancer cohorts in TCGA and ICGC (Figure 1C).

We calculated the mutation frequency for each gene within each cancer type, and highlighted the top 20 mutated genes according to the average mutation frequency across all cancer cohorts (Supplementary Figure 1). In TCGA, the 20 most frequently mutated genes are protein-coding genes. *TP53* was the most conspicuous one with an average mutation frequency of 37.80%, followed by *TTN* (33.32%) and *MUC16* (19.95%). Gene length can positively affect the mutation rate within a gene. Thus, we labeled the gene length and its rank among all human genes. A few of these prioritized genes may have stood out partly due to a large gene length. For example, *LRP1B* ranked number eight in overall mutation frequency (12.88%) and number nine in gene length; *CSMD1* ranked number 18 in overall all mutation frequency (9.47%) and number six in gene length. Eighteen subjects in TCGA had mutations in all the prioritized genes (Supplementary Figure 1A).

The 20 most frequently mutated genes in ICGC presented a similar picture as in TCGA (Supplementary Figure 1B). *TP53* again was crowned with the greatest average mutation frequency at 29.04%. One noticeable difference between ICGC and TCGA is TCGC included two non-coding genes in its priority list: *TTN-AS1* (22.06%) and *FLG-AS1* (10.81%). Both non-coding RNAs are the antisense of their respective sense genes. This could be a byproduct of ICGC’s unique vehicle of whole genome sequencing platform, as compared to TCGA’s exome sequencing. Ten subjects in ICGC had mutations in all the prioritized genes (Supplementary Figure 1B). For DepMap, the 20 most frequently mutated genes were all protein-coding genes. *TTN* had the greatest average mutation frequency at 65.31%, followed by *TP53* (61.82%) *MUC16* (44.88%). Seven cell lines had mutations in all the prioritized genes (Supplementary Figure 1C).

### Overall co-mutation description

As expounded in Methods, we sought to identify two levels of co-mutation pairs: co-mutated mutation pairs and co-mutated gene pairs. For the three consortiums (TCGA, ICGC, CCLE) we identified 30,841, 563,168, 1,286,266 co-mutations at gene level, respectively (Figure 2A). The large difference in the numbers of co-mutation identified among the three consortiums may reflect the difference in the total number of subjects and methods related to sequencing and mutation calling. The most frequent genes appearing in co-mutation pairs were identified (Figure 2B). For TCGA, the top genes were known cancer genes such as *TTN, MUC15*, and *PTEN*. For *ICGC*, the top gene was the non-coding gene *TTN*-*AS1*, followed by *MUC4* and *NBPF20*. For DepMap, the top genes were *TTN, MUC16* and *SYNE1*. Compared with the gene-level findings, co-mutations at position level have much lower frequencies, and accordingly we identified 0, 17, 63 co-mutations for TCGA, ICGC and DepMap, respectively (Figure 2C). All of the position level co-mutations with frequency ≥ 10% were contributed by ICGC’s thyroid carcinoma China (THCA-CN) and DepMap’s colon cohorts. The complete list of co-mutation pairs can be found in Supplementary Table S1 and S2 for the gene level and the position level, respectively.

**Figure 2.**
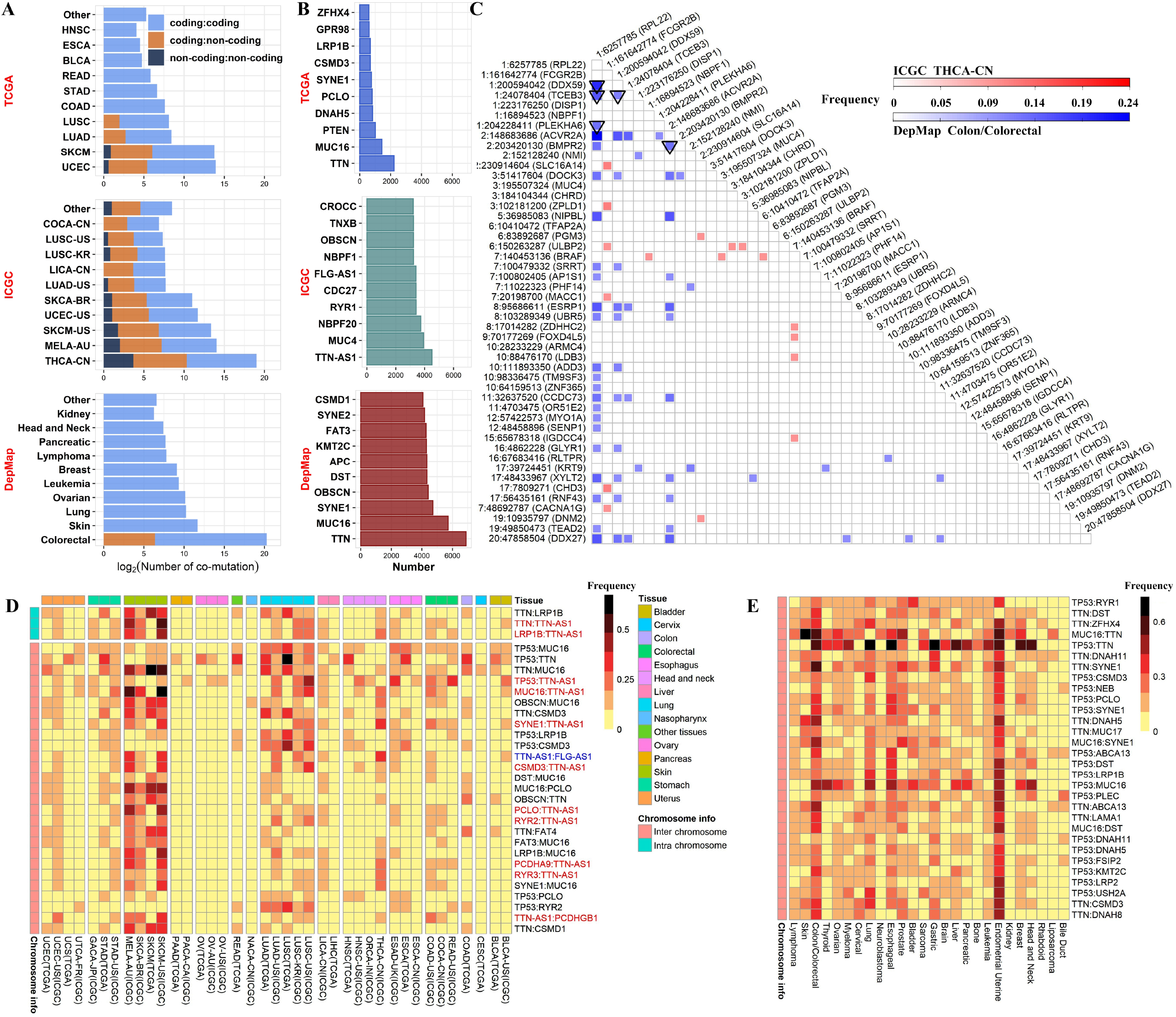
Overall co-mutation description. A. Amounts of co-mutation gene pairs identified from each cohort of three separate cancer consortiums. TCGA, The Cancer Genome Atlas. ICGC, International Cancer Genome Consortium. DepMap, Cancer Dependency Map. As indicated in the legend, three types of co-mutation pairs were distinguished: coding vs. coding, coding vs. non-coding, and non-coding vs. non-coding. B. Top ten genes most frequently appearing in co-mutation pairs. C. A total of 80 co-mutation position pairs were discovered, and they were indicated as the colored cells in the triangle heatmap identified by the row axis and column axis. Color scale is proportional to the frequency of a co-mutation pair. Red, originating from ICGC’s THCA-CN cohort; blue, originating from DepMap’s Colon cohort. Square, inter-chromosomal co-mutations; triangle, intra-chromosomal co-mutations. All genomic positions in panel C are based on GRCh37 human reference genome. D. The top 30 co-mutation gene pairs commonly shared across TCGA/ICGC cohorts. Co-mutation pairs involving a non-coding genes were distinguished in red font. E. The top 30 co-mutation gene pairs commonly shared across DepMap cell lines

Next, we examined co-mutation across multiple cancer types by computing the most commonly shared co-mutations. Because TCGA and ICGC were both based on cancer subjects and they shared a large portion of cancer types, these two data sources were combined into one round of analysis. The top 30 co-mutation gene pairs commonly shared across TCGA/ICGC cohorts are depicted in Figure 2D. Three co-mutations were intra-chromosome, and 27 were inter-chromosome. The top intra-chromosome co-mutation was *TTN*:*LRP1B*, which occurred in 16 of 39 cancer types. For inter-chromosome co-mutations, *TP53*:*MUC16* took the lead, which occurred in 25 of 39 cancer types. All top 30 commonly shared DepMap mutations are inter-chromosome co-mutations (Figure 2E), where *TP53*:*RYR1* stood out by occurring in 20 of 26 tissue types.

### Co-mutation disparity with age, sex, and race

Regression analyses were conducted to determine if co-mutations have associations with age, sex, and race (Supplementary Table S3). For age, 14,896 significant co-mutation associations were identified. Mutations are the natural products of aging, as evidenced by the fact that healthy senior subjects tend to accumulate more mutations than young controls [36]. An intuitive expectation might be that co-mutations in cancer patients are positively correlated with age. However, the association results between age and co-mutation status illustrated a rather striking image (Figure 3A). The majority of the significant associations were located in the UCEC cohorts of TCGA and ICGC. The association directions were very much cancer dependent, with UCEC showing predominant negative associations (TCGA UCEC 99.99% negative, ICGC UCEC 99.97% negative). Other cancer types had a few sporadic results with associations from both directions. Here, we demonstrate the age association with the co-mutation *TP53*:*IDH1* in TCGA’s low grade glioma (LGG) which had an adjusted p-value of 4.07 × 10^−9^ (Figure 3B). The subjects with wildtypes of co-mutation *TP53*:*IDH1* mostly had an older age than subjects with co-mutation *TP53*:*IDH1*. The same trend held for both *TP53* and *IDH1* when we examined the mutation in only one gene.

**Figure 3.**
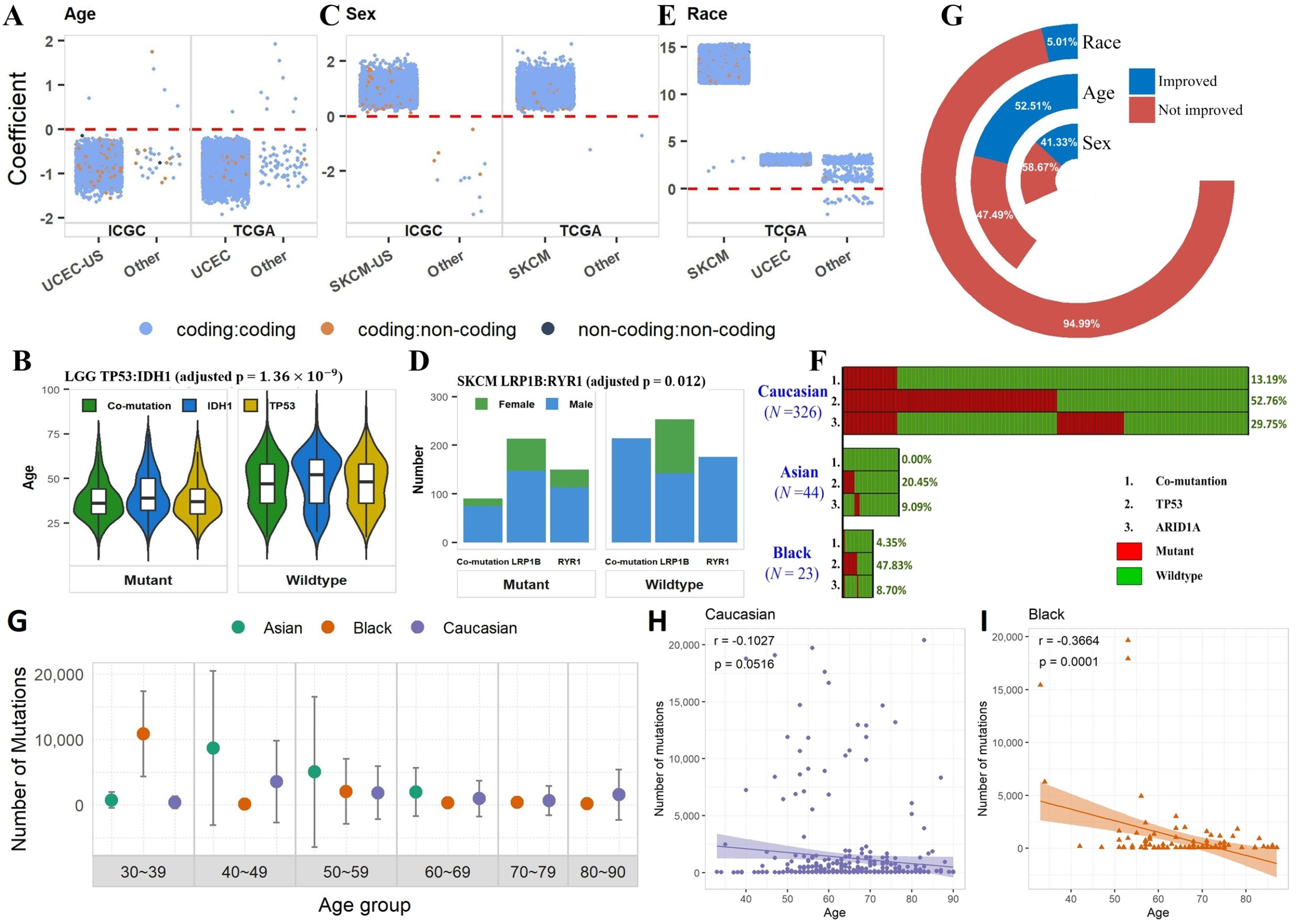
Association between co-mutation gene pairs and three phenotypic variables. A, association with age. C, association with sex. E, association with race. Each data-point in A, C, and E represents the regressed coefficient of one co-mutation with respect to a phenotypic variable. Only statistically significant co-mutations were plotted. The color of the dot represents the type of the co-mutation (coding:coding, coding:non-coding, and non-coding:non-coding). B, co-mutation pair *TP53*:*IDH1* demonstrated significant association with age in TCGA’s LGG cohort. D, co-mutation pair *LR1B*:*ZNF831* demonstrated significant association with sex in TCGA’s SKCM cohort. F, co-mutation pair *TP53*:*ARID1A* demonstrated significant association with race in TCGA’s BLCA cohort. F, Composition of phenotyp-associated co-mutation pairs in terms of improvement of association significance relative to single mutation association. G. Mutational burden in TCGA’s UCEC cohort by age group and race. H, I. Scatter plots of mutational burden vs age in TCGA’s UCEC cohort for Caucasian and Black. Pearson correlation coefficients and p-values were labeled on the scatter plots.

For sex, we found significant associations for 6,600 co-mutation gene pairs, most of which arose in the SKCM cohorts (Figure 3C). All significant co-mutations in SKCM showed a positive association in both TCGA and ICGC regardless of statistical significance, indicating males generally have a greater amount of co-mutation instances than females. If we ignore the statistical significance and examine the direction of associations for all co-mutations in skin-related cancers, we found that 98.9% of the 10,584 co-mutations in TCGA’s SKCM cohort had a positive association with sex; 99.1% of the 13,943 co-mutations in ICGC’s SKCM-US cohort had a positive association with sex; 98.3% of the 16,799 co-mutations in melanoma Australia cohort (MELA-AU); 81.1% of the 4,160 co-mutations in skin adenocarcinoma Brazil cohort (SKCA-BR) had a positive association with sex. These results suggest strong sex disparity for co-mutation and single mutation for skin cancer in general. Interestingly, significant associations between co-mutation and sex found in other cancer types (12 from ICGC’s STAD-US, LUSC-KR, and 2 from TCGA’s STAD, KIRC) indicated an opposite association trend, i.e., females having more co-mutations than males. Using the co-mutation *LRP1B:RYR1* as an example, the wildtype group consisted entirely of males, and the mutant group consisted 16.67% female. The deciding factor was the *LRP1B* gene with *LRP1B* mutant group contained 30.99% female and *ZNF831* wildtype group contained no female (Figure 3D). A few previous studies [37, 38] have shown the gender difference, with males showing higher mutations than females, as well as worse survival in male patients. They suggest that female melanoma patients have a statistically significantly higher frequency of tumor-associated, antigen-specific CD4+ T-cells than their male counterparts. This may lead to a more robust anti-tumor immune response in female patients that eliminates cancer cells even when they only have a small number of mutations and they cannot accumulate high mutations. As a result, female patients may have fewer mutations on average and better survival.

Finally, for race, 27,726 significant associations were detected, with a majority found in TCGA’s UCEC and SKCM cohorts (Figure 3E). We demonstrate the race disparity with the co-mutation *TP53*:*ARID1A* from TCGA’s BLCA cohort as an example. This co-mutation had a 13.19% frequency in Caucasian, 4.35% in Black, and 0.00% in Asian (Figure 3F). For Asian subjects, *TP53* had a frequency of 20.45% and *ARID1A* had a frequency of 9.09%. Yet, the two mutant groups did not overlap on a single subject. Furthermore, of the significant associations for age, sex, and race, 52.5%, 41.3%, and 5.0%, respectively, had stronger effects for co-mutation than their corresponding single mutations (Figure 3G).

The above results suggest that age, sex, and race play significant roles in co-mutation and possibly single mutation as well. TCGA’s UCEC cohort is a unique example. Dividing UCEC subjects into age and race groups, younger subjects tend to have higher mutational burdens (Figure 3G). The negative correlation between mutational burden and age is marginally significant for Caucasians (Figure 3H) and significant for Black (Figure 3I) and not significant for Asians which may be due to limited sample size.

### Survival Analysis

At position level, we identified 17 co-mutations from ICGC data and none from TCGA data with frequency ≥ 10%. All 17 position-level co-mutations came from ICGC’s THCA-CN cohort, where no events were recorded at the time of data collection. Thus we were unable to conduct any survival analysis at position level. At gene level, after multiple-test correction, eight co-mutations were found to be significantly associated with survival (adjusted p-value < 0.05, Table 1, Figure 4A). Five of the eight were from TCGA and three were from ICGC. The most significant one was the co-mutation *TP53*:*KRAS* in ICGC’s pancreatic cancer cohort (PAAD-US) (HR = 2.87, 95% CI 1.71-4.84). The second most significant co-mutation, *TP53*:*TTN-AS1*, involves a non-coding gene, and it was mined from ICGC’s ovarian cancer Australia cohort (OV-AU) (HR = 2.16, 95% CI 1.22-3.85). The co-mutation *TP53*:*KRAS* in TCGA’s PAAD cohort also achieved a significant association (HR = 1.91, 95% CI 1.21-3.05), ranked in the 6^th^ place overall by adjusted p-value. To demonstrate that some co-mutations can have a better prognosis value than their corresponding single mutations, we conducted single mutation survival analyses, where the subjects were divided based on mutation status of a single gene. Of the eight significant co-mutations, three had a more significant p-value than both of their corresponding single mutations. These three included *TP53*:*ATRX* in TCGA’s LGG cohort, *TP53*:*KRAS* in ICGC’s PAAD cohort, and *KMT2D*:*BCL2* in ICGC’s German malignant lymphoma cohort (MALY-DE).

**Figure 4.**
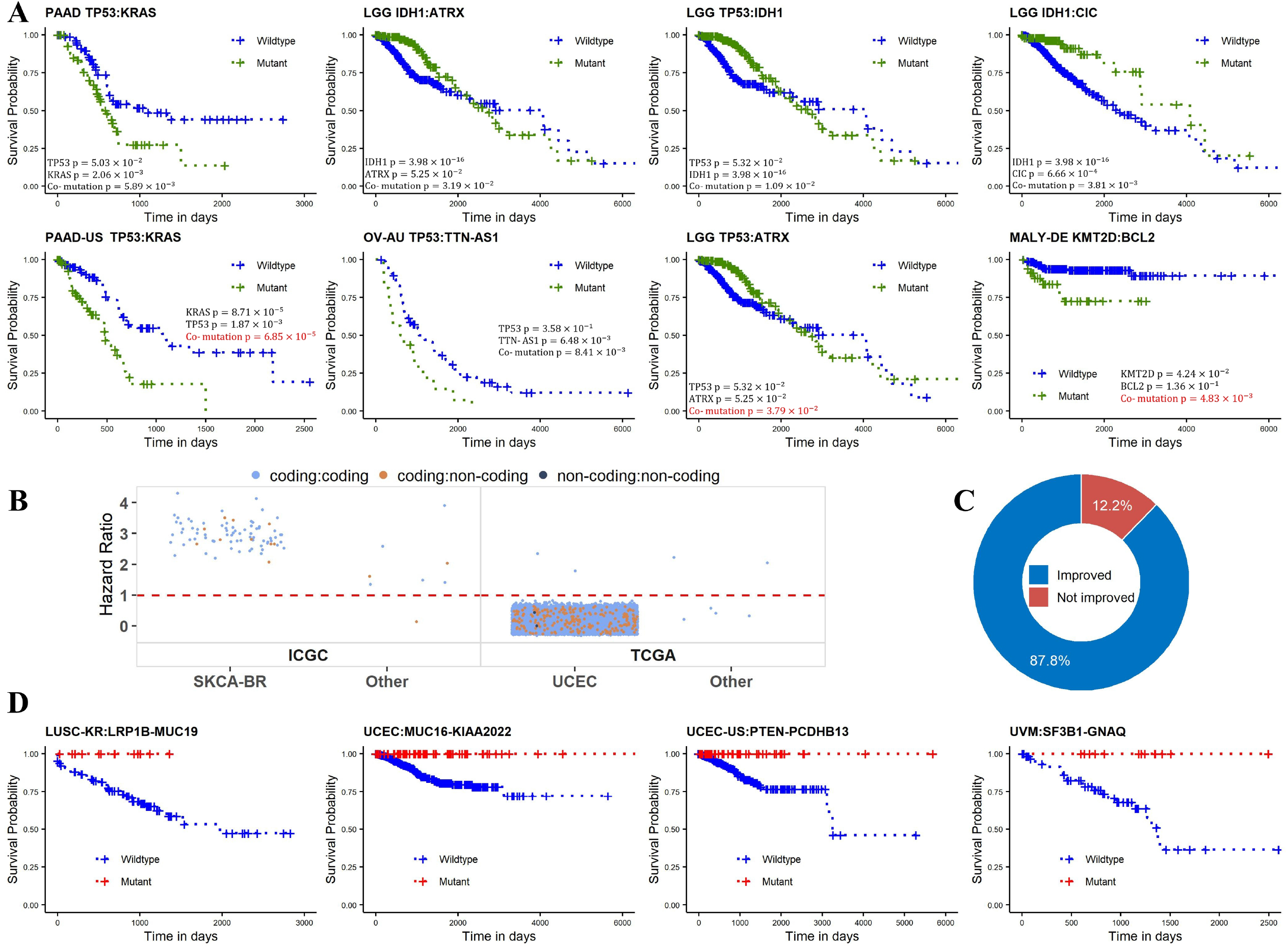
Prognosis power of co-mutation gene pairs for cancer patients. A. Kaplan-Meier curves of eight prognostic co-mutation gene pairs as inferred from the Cox-proportional hazard regression model. Unadjusted p-values for the co-mutation analysis and the two single-gene analyses, all using the Cox-proportional hazard regression model, were labelled in each Kaplan-Meier plot. A p-values in red font highlights the scenario where the co-mutation p-value was more significant than the respective single-gene analysis p-values. B. Directional analysis of marginally significant co-mutations (0.05 < adjusted p-value < 0.1). In ICGC SKCA-BR cohort, all 82 marginally significant co-mutations had an HR great than one. In TCGA’s UCEC cohort, 14,350 of 14,352 marginally significant co-mutations had an HR less than one. C. Composition of prognostic co-mutation pairs in terms of improvement of prognosis power relative to single-mutation power. D. Four representative prognostic co-mutations ascertained due to imbalanced events. All death events occurred in the wildtype groups.

Furthermore, 14,440 co-mutations were found to be marginally significant (0.05 < adjusted p-value < 0.1) (Supplementary Table S4, Figure 4B). Of the 14,440 marginally significant co-mutations, 87 were from ICGC, including 82 from the SKCA-BR. Of the 14,353 marginally significant co-mutations from TCGA, only one (*TP53*:*DNAH5*, HR = 2.00, 95% CI 1.26-3.17) was from the head and neck squamous cell carcinoma (HNSC) cohort, and all the rest ones came from UCEC. Similar to the sex disparity of co-mutation association observed earlier, the direction of survival prediction was remarkably cancer dependent. In SKCA-BR cohort, all 82 co-mutations had HR greater than one, indicating better prognosis for the wildtype groups. In TCGA’s UCEC cohort, of the 14,352 marginally significant co-mutations, 14,350 (99.9%) had HR less than one, indicating better prognosis for the mutant groups.

When conducting the Cox proportional hazard regression model, there is a scenario that either the mutant or the wildtype group did not have any event. As explained in the method section, we termed this group of co-mutation as significant due to imbalanced events. A total of 246 such co-mutations were identified, 226 were from TCGA’s UCEC cohort (Supplementary Table S5). All of these 246 co-mutations favored better prognosis, meaning that the co-mutation mutant groups did not report any death event, and the wildtype group had at least 10 death events. To demonstrate whether co-mutations can provide a better prognosis than single mutations, we counted the survival events within single-gene-mutant groups. If single-gene-mutant groups for both constituent genes of the pair had non-zero death events, we concluded that the co-mutation provided additional prognostic value than both corresponding single mutations. Of the 246 co-mutations, 216 had improved prognostic value than single mutations (Figure 4C). In another word, if we had divided the subjects into mutant and wildtype groups based on single mutations, the scenarios of imbalanced event distribution would not have occurred. This demonstrated that additional prognostic power was offered by the co-mutation gene pair as compared to the corresponding single gene mutations. Kaplan-Meier curves for four example co-mutations from these 246 are displayed in Figure 4D. Because no event occurred for the mutant group, the mutant probability trends came out as flat lines. Since sex disparity in co-mutation frequency was demonstrated earlier, we also conducted survival analysis based on sex. TCGA’s glioblastoma (GBM) cohort had the most significant result (HR:1.44), but it did not pass multiple test correction.

### Functional analysis

Survival association may be an indication for functional variants. The eight prognostic co-mutations with adjusted p-value < 0.05 and the 254 prognostic co-mutations with empirical significance due to imbalanced events involved 144 distinct genes altogether. Using the mutations in these 144 genes, we conducted somatic binding sequence analysis to determine if these mutations caused any alteration in TF, RBP, miRNA seed, and miRNA-matching 3’-UTR binding sequences. The analyses revealed 13,192 gains and 12,969 losses in RBP binding sequences (Figure 5A, Supplementary Table S6) and 5,830 alterations in miRNA-matching 3’-UTR binding sequences (Supplementary Table S7). In total, we found mutations of 131 genes resided in RBP binding sequences, and mutations of 121 genes resided in miRNA-matching 3’-UTR binding sequences. For example, *TP53*, one gene frequently appearing in co-mutation pairs, had a mutation (C→T) at chromosome 17 position 7,676,273 (GRCh38 coordinate) in TCGA’s rectum adenocarcinoma cohort (READ). This mutation caused losses of binding sequences for RBPs SRSF1 and SRSF2, but also rendered gains of binding sequences for two RBPs RBMX and SRSF3. The *SRSF* gene family encodes for the serine and arginine rich splicing factors. Genes of this family has been frequently associated with cancers [39-41]. RBMX is a chromosome-x-linked RNA binding motif protein, which has also been associated with bladder cancer [42] and kidney cancer [43]. A good example for altered miRNA-matching 3’-UTR binding sequences is the *TP53* mutation (C→T) at chromosome 17 position 7,673,780, which altered the binding sequence to *miR-150-5p* in TCGA’s LGG cohort. The miRNA *miR-150-5p* has been found to suppresses tumor progression by targeting VEGFA in colon cancer [44]. This altered binding sequence can potentially disrupt the normal regulation between *TP53* and *miR-150-5p*. From these two example mutations in *TP53*, we show the intricate consequences of mutations. The combinatorial effect arising from concurrent mutations will further complicate the disruption of binding sequences.

**Figure 5.**
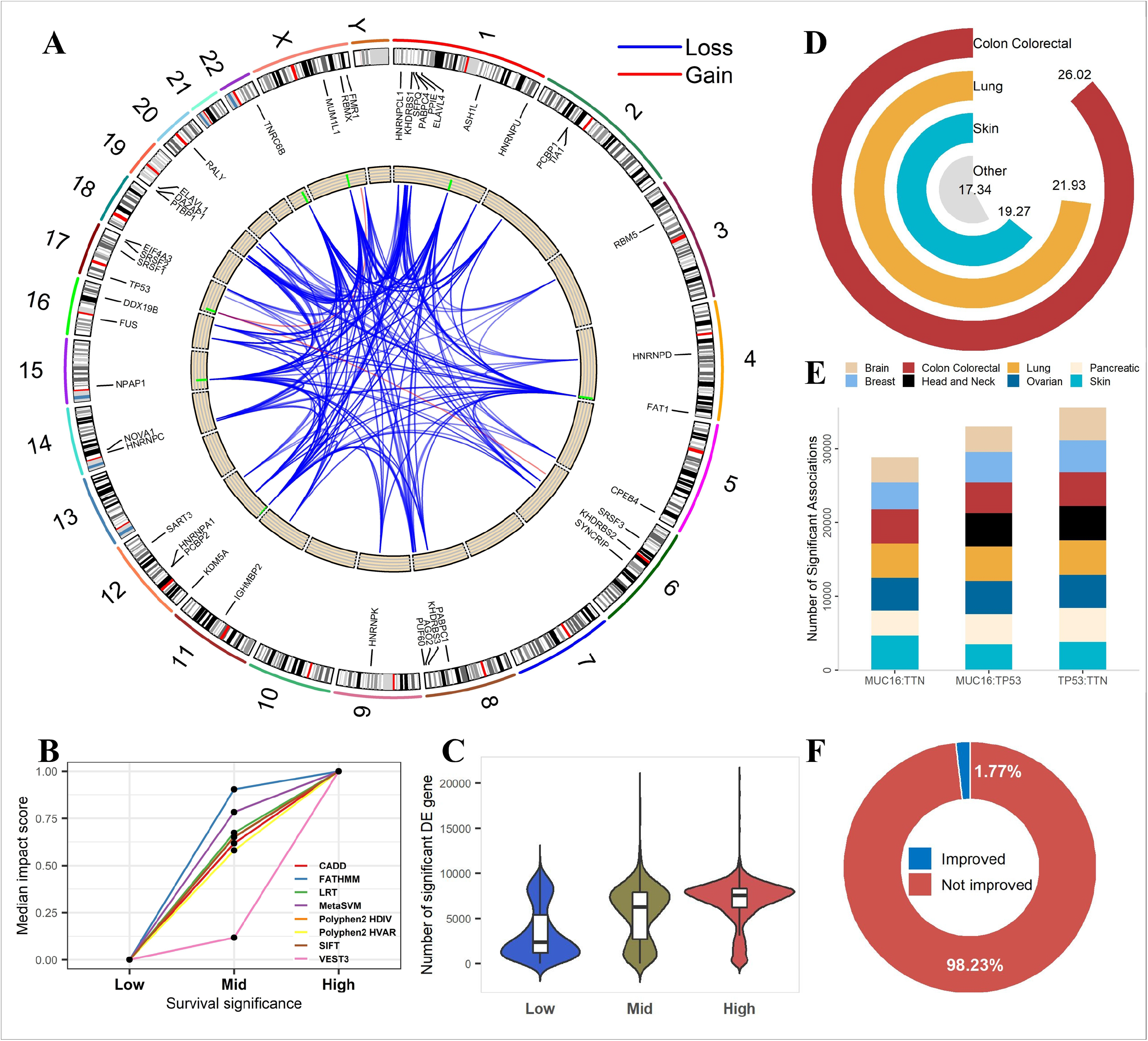
Functional characterization of genes involved in co-mutation pairs. A. A total of 26,161 alterations in RBP binding sequences were attributed to the prognostic concurrent mutations (significant co-mutations resulting from the survival analyses). Using ⩾ 1% as the threshold, we further filtered 6 gains (red) and 965 losses (blue) of RBP binding sequences to plot. The ends of lines in the middle circle represent the position of mutation and its affected RBP. The green bars in the inner ring represent the frequency of the mutation. B. Average mutation impact scores of three prognostic-level co-mutation groups. Mutation impact scores were predicted by eight algorithms that were specified in the legend. C. Amounts of significantly differentially expressed genes between the mutant subjects and the wildtype subjects, as determined by the co-mutation status. Like in B, three prognostic-level co-mutation groups were analyzed separately and compared between each other. D. A donut plot to show the (log2-scaled) numbers of significant associations between co-mutation and drug sensitivity. E. The top three co-mutations with the most significant drug sensitivity associations. F. Composition of drug-sensitivity associations for co-mutation pair *TP53*:*KRAS* in terms of improvement of co-mutation significance over the single-mutation significance.

We also conducted several functional predictive analyses using eight established prediction algorithms. From all significant co-mutations from the above survival analysis, we obtained three groups with each containing a distinct set of 1000 co-mutations. The three groups represented co-mutations at the bottom, medium, and top level, respectively, based on the p-value out of the Cox model analysis. The median impact scores for these three groups were plotted (Figure 5B). All eight mutation impact prediction scores produced consistent results, co-mutations that were more significantly associated with survival tend to have stronger impact scores.

In addition to the theoretical prediction analyses, we also conducted empirical data based impact analysis. We conducted differential gene expression analysis between co-mutation mutant and wildtype groups. Across the bottom, medium, and top survival association groups, we compared the numbers of differentially expressed genes (adjusted p-value < 0.05). As demonstrated in Figure 5C, for co-mutations, the number of differentially expressed genes increased with the survival significance level.

Furthermore, we computed co-mutation’s association with drug sensitivity using data from DepMap consortium. We tested 23,486 co-mutations and sensitivity from 4,686 drugs. Overall, we detected 72,639,066 significant associations (adjusted p-value < 0.05) (Figure 5D, Supplementary Table S8). The majority of the co-mutations of significant drug sensitivity were contributed by the colon cancer cell lines. The three co-mutations most frequently involved in significant drug sensitivity associations were *TP53*:*TNN, MUC16*:*TP53*, and *MUC16*:*TNN* which had 35,621, 33,012, and 28,871 significant drug sensitivity associations, respectively (Figure 5E).

Moreover, we used the co-mutation *TP53*:*KRAS* as an example to prove that a co-mutation may provide additional information than the single mutations entailed therein. This co-mutation was found to be significantly associated with survival in both PAAD cohorts in TCGA and ICGC. Drug sensitivity analysis found 4,684 significant associations for co-mutation *TP53*:*KRAS*, of which 83 had co-mutation p-values more significant than the corresponding single mutations’ p-values (Figure 5F).

### Comparisons with clinical cancer gene panels

Currently, a majority of hospitals test cancer patient biopsy with an established cancer gene panel to guide the treatment strategy. The four panels, namely Agilent SureSelect, University of California San Francisco UCSF500, FoundationOne CDx, and Ashion Genomic Enabled Medicine, contained a total of 898 distinct cancer genes. We compared these four panels with our survival significant co-mutation genes. Ignoring marginally significant prognostic co-mutations, we considered the eight rigorously significant co-mutations and the 246 significant co-mutation due to imbalanced events, which involved a total of 144 genes. An intersection examination found that there is a large disagreement among the cancer panels (Supplementary Figure 2). There are 72 common genes across the four cancer panels. Of the 144 co-mutation genes, 38 are covered by the four cancer panels, 106 are not covered in any of the four panels. Our analysis results have shown that many of these 144 co-mutation genes have potential functional impact and prognostic value. Adding these co-mutation genes to the cancer panel may be beneficial to cancer patients, because they help to provide a more accurate description of impactable mutations, and offer potential alternative treatment plans.

## Discussion

Accumulation of somatic mutations, especially driver mutations throughout life can lead to tumorigenesis. While the majority of the somatic mutation studies have been focused on single mutations, gradually, the importance of co-mutations has been established. Utilizing 29,559 cancer subjects and 1,747 cancer cell lines covering 78 distinct cancer types, we conducted the most comprehensive co-mutation study to date, uncovering several novel co-mutation related findings. The mutation data from the three consortiums provided an excellent overview of the landscape of co-mutations in cancer. The mutation spectrums can be different among the three consortiums due to the nature of the sample, sequencing type, and mutation calling method. The most noticeable difference is the number of mutations detected, which is much higher for DepMap. We speculate that this is because cell lines were cultured from a single cell of tumor which allow easier identification of mutations. Even with the difference, some patterns were blatantly visible across all three consortiums. For example, the bimodality of mutated genes for colon cancer can be seen across all three consortiums.

One of the interesting findings is related to the sex disparity of co-mutation in skin cancers. The sex disparity of single mutation for skin cancer has been discussed by a previous study [37], in which the authors also demonstrated sex disparity in TCGA’s SKCM cohort. The author mentioned that one of the limitations of the TCGA data is the exome sequencing which only allowed the detection of sex disparity in exome regions. In our analysis, ICGC’s MELA-AU and SKCA-BR cohorts were with whole genome sequencing and also displayed a strong disparity favoring more co-mutations for males. Our results reinforced the finding of single mutation sex disparity in skin cancer and demonstrated that such disparity can be expanded to co-mutations.

One of the major goals of our study is to show that co-mutations provide additional information compared to their corresponding single mutations. The advantage of co-mutation was primarily demonstrated through our survival analysis, in which we identified eight co-mutation that were significantly associated with survival. And three of the eight co-mutations provided better prognostic prediction than their corresponding single mutations. The same concept was then again demonstrated in 216 of 246 significant co-mutations that did not have events in the mutant groups. More strikingly, our results uncover cancer dependent survival association directionality. For ICGC’s SKCA-BR cohort, all 82 marginally significant co-mutations had HR greater than one, suggesting better prognosis for the wildtype groups. In contrast, TCGA’s UCEC cohort had 14,352 marginally significant co-mutations, and 99.9% had HR smaller than one, indicating poor prognosis for the wildtype groups. The phenomenon of higher mutational burden is beneficial for survival has been observed in metastatic melanoma [37] and patients with higher mutational burden responded better in a trial of Ipilimumab [45]. However, the same phenomenon has not been reported in uterine cancer. The survival association for TCGA UCEC’s mutational burden was marginally significant (HR: 0.9998, 95% CI (0.9997–1), p-value = 0.05). The direction of HR indicates higher mutational burden is better for survival. This may suggest a similar mechanism between melanoma and uterine cancer.

Certain mutations when occurred simultaneously can produce stronger tumorigenesis or protective effect, which can translate to better prognostic prediction. In certain cancers, the directions of co-mutation survival are remarkably consistent, which suggests cancer dependent mutation mechanisms. From our analysis, skin cancer and uterine endometrial cancers frequently showed up as cancer types with extreme results. Our analysis demonstrated that the uterine endometrial cancer subject’s mutational burden is negatively correlated with age. This is consistent with uterine cancer’s etiology which can be classified into two categories by age: 1) for younger pre-menstrual women, endometrial cancer usually occurs with excessive endometrial growth, and the secretion of excess estrogen can not be balanced with progesterone; 2) for older post-menstrual women, cancers are not caused by the high level of estrogen secretion [46]. We speculate that this may be due to the hypermutated subjects within these cancer types. And these co-mutations may be representing overall cancer specific mechanisms because of the consistencies of observation for all co-mutations in these cancer types.

After determining co-mutation’s prognostic value, we examined the potential functional impact of co-mutations theoretically and empirically. Theoretically, we used eight mutational impact prediction tools to predict co-mutation’s overall impact. This analysis showed that co-mutations with more significant survival associations had higher impact predictions, suggesting the survival associations have potentially resulted from the functional impacts. Empirically, we examined co-mutation related gene expression dysregulation and drug sensitivity alteration.

The importantance of non-coding genes has been increasingly acknowledged. For example, an recent study found that the overall prognostic power increases with the addition of non-coding gene expression [47]. Our study focused on protein-coding genes mostly due to the limitation of data. However, a small percentage of relevant coding:non-coding and non-coding:non-coding co-mutations were also detected. For example, non-coding RNA *TNN-AS1* was detected in two co-mutations that were significantly associated with survival. With additional whole genome sequence data release in the future, we expect more impactful non-coding co-mutations can be identified.

From the clinical aspect, we showed that current cancer gene panels disagree and are missing many co-mutation genes we have discovered in this study. While we encourage the addition of the co-mutation genes into the cancer panels, we also acknowledge that whether each co-mutation is actionable requires further mechanistic study.

## Authors’ Contributions

L. Jiang, H. Yu conducted formal analysis. Y. Guo, S. Ness, P. Mao and L. Jiang wrote the manuscript. J. Tang, Y. Guo, and F. Guo supervised the study and provided funding.

## Acknowledgments

This study was support by Cancer Center Support Grant P30CA118100. This study was supported by Analytical and Translational Genomics Shared Resource and Bioinformatics Shared Resource of the Comprehensive Cancer Center, University of New Mexico. Y. Guo was supported by grant R01ES030993-01A1 from the National Cancer Institute, USA.

---

**Supplementary Figure 1.**
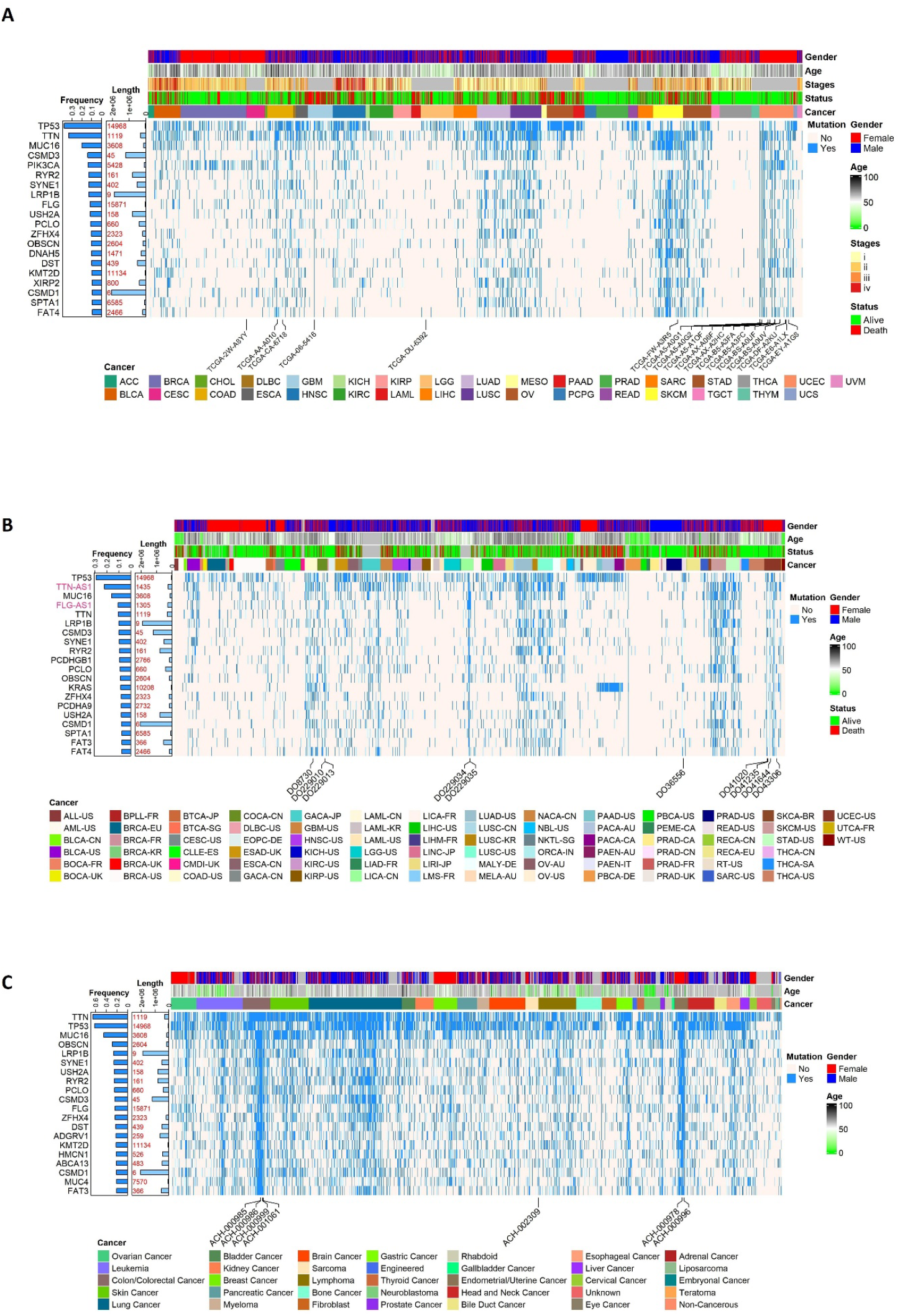
The twenty most frequently mutated genes in the three cancer data consortiums separately. A. The Cancer Genome Atlas (TCGA). B. International Cancer Genome Consortium (ICGC). C. Cancer Dependency Map (DepMap). Each row of the heatmap represents a gene, and each column represents a subject showing mutation in at least one gene. Gene names in black denote protein-coding genes, and gene names in pink denote non-coding genes. Two barplots are attached at the left side of the heatmap to annotate two distinct features of the mutated genes. One barplot denotes the mutation frequency of each gene across all cohorts of one same consortium; the other barplot visualizes the gene length with the corresponding rank specified. Available phenotypic variables of subjects were indicated with color bars on the top of the heatmap. Subjects bearing mutations in all 20 genes are identified below the heatmap.

**Supplementary Figure 2.**
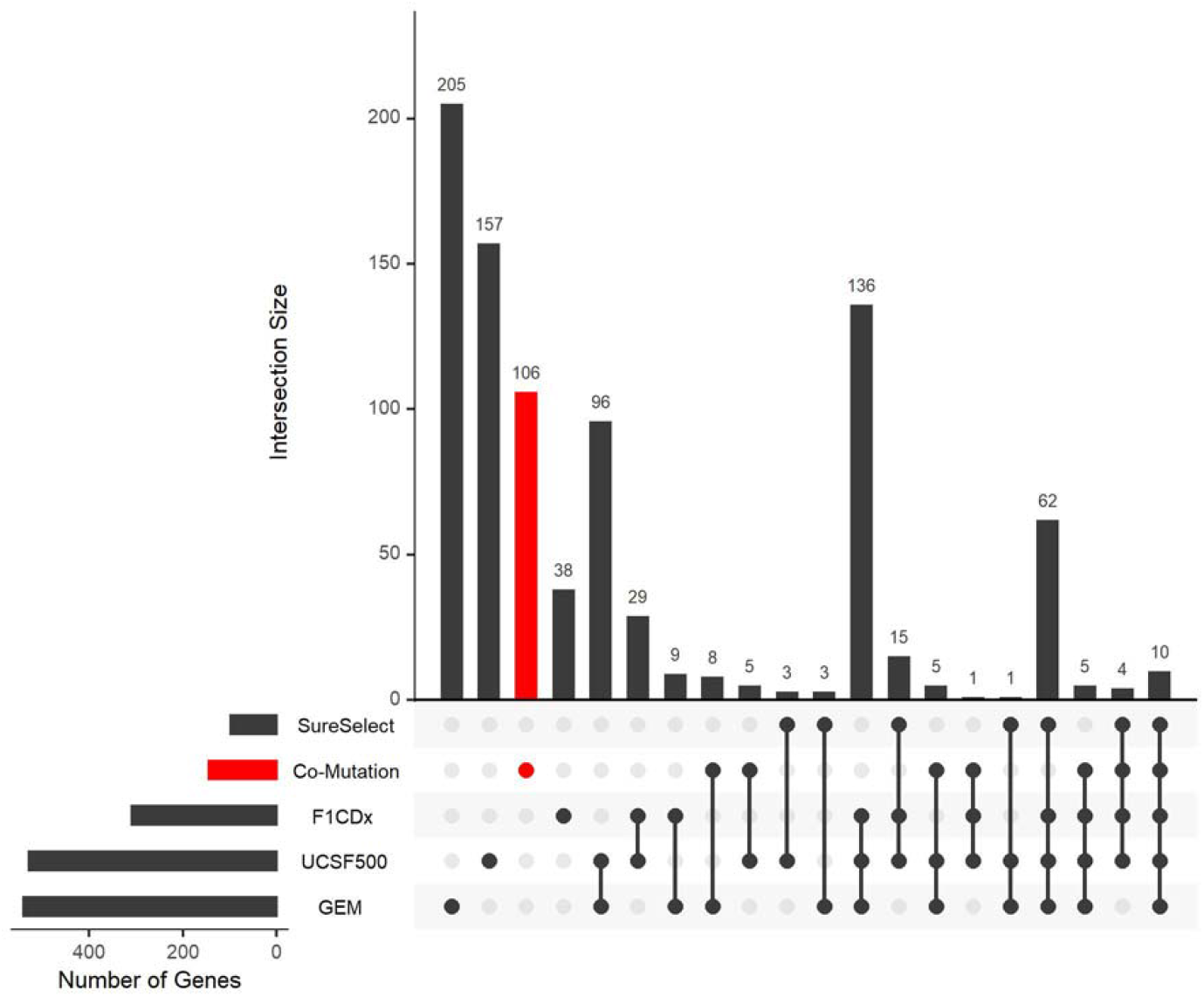
Intersection analysis plot among five different cancer-related gene sets. The five gene sets included our identified co-mutation genes (red) and four clinical cancer gene panels (black), whose identification and gene number are depicted as horizontal barplot on the bottom left panel. The unitary, binary, tertiary, quaternary, and quinary intersection relations were illustrated with line segments on the bottom panel. The actual size of each intersection set formed by one, two, three, four, or five sets out of the total five was depicted in the vertical barplot on the top main panel. For example, the first verticle bar (205), represents the unique genes in GEM cancer gene panel and these 205 genes are not included in the other four gene sets. The last column (10) represents the overall intersection of the five gene sets. The fourth to the last column (62), represents the intersection of the four cancer panels minus co-mutation gene sets. Thus, the four cancer gene panels share 62 + 10 = 72 genes. SureSelect, Agilent SureSelect. F1CDx, FoundationOne CDx. UCSF500, University of California San Francisco UCSF500. GEM, Ashion Genomic Enabled Medicine.

